# Heartbeat-locked auditory deviations slow down cardiac activity

**DOI:** 10.64898/2026.04.28.721302

**Authors:** Matthieu Koroma, Kevin Nguy, Andria Pelentritou, Marzia De Lucia, Athena Demertzi

**Affiliations:** Physiology of Cognition Lab, GIGA-CRC Human Imaging Unit, GIGA Institute, Allée du 6 Août, 8 (B30), 4000 Sart Tilman, University of Liège, BELGIUM; Fund for Scientific Research FNRS, Brussels, BELGIUM; Psychology & Neuroscience of Cognition (PsyNCog), Place des Orateurs, 1 (B33), 4000 Sart Tilman, University of Liège, BELGIUM; Brain-Body and Consciousness Laboratory, Lausanne University Hospital (CHUV) & University of Lausanne, Lausanne, Switzerland

**Author notes:** **Corresponding author**: Athena Demertzi, PhD, Physiology of Cognition Lab, GIGA-CRC Human Imaging Unit, Allée du 6 Août, 8 (B30), 4000 Sart Tilman, University of Liège, BELGIUM.

**Keywords:** interoceptive-exteroceptive integration, cardio-audio synchrony, ECG, cardiac surprise, active inference, dynamical coupling

## Abstract

Our responses to environmental inputs depend on the variations of our own physiological activity. However, the mechanisms by which the integration of sensory information with interoceptive signals shape bodily responses to external events remain debated. In this pre-registered study, we hypothesized three possible mechanisms underlying such exteroceptive-interoceptive integration: cardiac surprise, active inference, and dynamic coupling. To test them, we implemented a closed-loop stimulation procedure to play auditory deviations from sequences either synchronized or not with heartbeats which varied in type (omissions or rare tones) and predictability (random or regular intervals). First, we replicated previous findings that cardiac activity slows down in response to sound omissions only when sounds are synchronized with heartbeats. Second, we showed that this effect extends to rare tones, excluding the dynamic coupling hypothesis. Third, we demonstrated that these responses do not depend on the predictability of auditory deviations, excluding both cardiac surprise and active inference hypotheses. In a control experiment, we further observed that behavioral responses depend on the type and predictability of auditory deviants: participants can discriminate subjectively which sounds were synchronized with their own heartbeats without evidence of a relationship to interoception nor cardiac responses. Overall, these results demonstrate that auditory deviations slow down cardiac responses when locked to heartbeats but independently from their type and regularity, calling for novel hypotheses to account for the interoceptive-exteroceptive integration of sensory signals into cardiac activity.

**Impact statement:** Using a cardio-audio synchrony task, we show that cardiac responses slow down upon heartbeat-locked auditory deviations independently from their type or regularity, suggesting a simple, fundamental mechanism of integration of internal and external signals into bodily responses to the environment. The heart may provide a straightforward way to study basic self-related processes, without depending on behavior or self-report, which is especially valuable for individuals who are unable to respond or communicate.

## Introduction

Bodily information influences our perception and ability to react to the environment (Azzalini et al., 2019; Hsueh et al., 2023). More specifically, cardiac signals play an important role in emotional processes (Garfinkel et al., 2014; Hsueh et al., 2023; Pfeifer et al., 2017), self-consciousness (Seth & Critchley, 2013; Tallon-Baudry et al., 2018), social perception (Azevedo et al., 2017) and perceptual processes (Al et al., 2020; Arslanova et al., 2023). For example, the relative timing of heartbeats to sensory stimuli impacts stimulus recognition (Sandman et al., 1977) and detection (Saxon, 1970), while the timing and type of sensory stimulation was shown to modulate cardiac rhythms (Pérez et al., 2021; Raimondo et al., 2017; Koroma et al., 2025). The exact nature of mechanisms supporting the bidirectional interaction between sensory stimuli and visceral signals remains nevertheless debated (Palmer & Demos, 2022; Skora et al., 2022).

Several models have been proposed to explain such interoceptive-exteroceptive integration. An influential theory accounting for the interaction of bodily activity and perception is interoceptive predictive coding (Seth, 2013; Seth & Critchley, 2013). According to this framework, visceral signals are dynamically updated based on the organism’s own prior predictions to reduce prediction errors arising from the comparison with incoming sensory inputs. This mechanism can explain how unpredicted events influence heart rate to prepare the body for potential threat detection (Ekman et al., 1983; McCorry, 2007). This cascade leads to faster and more accurate perceptual detection (Anllo-Vento, 1995; Mangun, 1995), lower detection thresholds (Correa et al., 2005; Hawkins et al., 1990; Luck et al., 1994) and enhanced computation of temporal regularities in sensory processing (SanMiguel et al., 2013).

Active inference, a related account stemming from predictive processing, puts a stronger focus on how organisms use both internal and external activity to minimize prediction errors (Pezzulo & al., 2014). Unpredicted mismatches between interoceptive and sensory activity can be minimized by descending visceral control, *e*.*g*., vagal activity, explaining how heart rate adapts to the occurrence and predictability of external stimuli to maintain optimal neural and bodily efficiency (Seth & Friston, 2016; Seligowski et al., 2021).

Finally, dynamical coupling emphasizes that these processes occur within a continuous organism–environment loop and posits that internal oscillatory systems are synchronized to external inputs, allowing to reduce metabolic costs in perceptual processing and enhance behavioral performance (Greenfield et al., 2021; Lakatos et al., 2019; Lim et al., 2014; Terry et al., 2012). According to this model, heart rate aligns with the timing of sensory inputs without the need to resort to predictions (Palmer & Demos, 2022).

To investigate these mechanistic accounts of the integration of visceral with external sensory processing, we relied on the cardio-audio synchrony paradigm that uses a closed-loop auditory stimulation to present sound sequences, either at a short-fixed delay after heartbeats, *i*.*e*., synchronously, or without any systematic relation to heartbeats, *i*.*e*., asynchronously (Pfeiffer & De Lucia, 2017).

Building upon previous evidence that sound omissions presented synchronously to heartbeats slow down cardiac activity (Pelentritou et al., 2024, 2025), we designed a pre-registered protocol (Koroma et al., 2023) that probes cardiac responses to auditory deviants within sequences locked to the participant’s heartbeat or not. Sensory deviations varying in type, including either omissions or rare tones, were played with different predictabilities, being presented at either regular or random intervals.

We hypothesized that interoceptive-exteroceptive integration during cardio-audio synchrony would lead to slower cardiac responses to deviants when sound sequences were synchronized with heartbeats. With respect to the tested mechanisms, we expected: i) in the cardiac surprise model: a main effect of predictability, irrespective of the type of sensory stimulation, because deviants would be less surprising when predictable, ii) in the active inference model: an interaction effect between the type and regularity of sensory stimulation, because cardiac responses would be adapted to stimuli only when they could be predicted, iii) in the dynamical coupling model: a main effect of deviant type, irrespective of the regularity of deviants, because responses would be aligned to external signals independent from their predictability (Table 1).

**Table 1.** Pre-registered hypotheses for cardiac responses to deviants within the cardio-audio closed-loop stimulation paradigm based on the proposed models of interoceptive-exteroceptive integration of cardiac and auditory signals. >: cardiac deceleration; = : no modulation.

|  | **Models** |  |  |
| --- | --- | --- | --- |
| **Conditions** | *Cardiac Surprise* | *Active Inference* | *Dynamical Coupling* |
| *Cardio-audio Synchrony* | synchronous > asynchronous | synchronous > asynchronous | synchronous > asynchronous |
| *Deviant's type* | omission = rare tone | omission > rare tone | omission > rare tone |
| *Predictability* | random > regular | random > regular | regular = random |

In addition, in a separate control experiment using the same design, we investigated whether these effects would have a behavioral consequence. According to our pre-registered hypotheses, we expected faster reactions to auditory deviants in synchronous over asynchronous conditions to account for the facilitatory effect of interoceptive-exteroceptive integration on the detection of auditory deviants. We also expect faster reactions as well as for rare tones over omission due to their increased saliency and for regular over random conditions due to their predictability.

As previous studies have highlighted that subjective awareness of the cardio-audio synchrony related to how accurately one can sense internal signals from their own body (Banellis & Cruse, 2021; Herbert et al., 2007; Sel et al., 2016; Suzuki et al., 2013; Tsakiris et al., 2011), we also asked participants across both experiments to report which sound sequences were time-locked or not to their heartbeats and investigated their interoceptive sensitivity using the Multidimensional Assessment of Interoceptive Awareness (MAIA) questionnaire (Willem et al., 2022).

We expected participants to be able to detect cardio-audio synchrony with a positive correlation with MAIA scores if they used interoceptive abilities. Similarly, we tested if cardiac responses to auditory deviants correlated with subjective responses and MAIA scores to study the relationship between cardiac signals, subjective awareness of heartbeat-locked auditory sequences and participant’s interoceptive sensitivity.

## Materials and Methods

### Procedure

The sample size was computed using Gpower, resulting in 34 participants for the main and 8 participants for the control experiment (within factors ANOVA, power: 0.95, middle effect: .25, 1 and 50 repeated measures for respectively the main and control experiment). To account for the risk of attrition in the number of participants, we independently recruited 40 and 10 participants for the main and control experiments, respectively, aged between 18 and 35 years old and equally balanced in sex and gender. All recruited participants were without a history of cardiac, mental disorders or auditory deficits. They provided written informed consent prior to participation and were free to abort the experiment at any time without facing negative consequences.

At the beginning of each experiment, participants were equipped with electrocardiography (ECG) amplified with the actiCHamp system (Brain Products, GmbH) to record cardiac responses. Behavioral responses were additionally recorded by button press in the control experiment. Across both experiments, subjective responses and questionnaires were collected with verbal and written reports. Finally, electroencephalography (EEG) (easy-cap 64 electrodes, actiCHamp amplifier), pupil (Drowsimeter R100; Phasya, S.A), electrodermal activity and respiration (MP160 amplifier; BIOPAC SYSTEMS Inc.) were also collected during the main experiment.

Upon completion of the main experiment, participants received a monetary compensation of 50 euros. In line with the Regulation (EU) 2016/679 of the European Parliament and Council from April 27th, 2016, data obtained during the experiment were anonymized and handled confidentially. The code of conduct aligned with ethical standards from the 1964 Declaration of Helsinki. The protocol was approved by the Ethics Committee of the University Hospital Liège (Nr. EudraCT : B7072022000046).

### Protocol

Stimuli were 50ms-duration pure tones sampled at 44100Hz generated with the Sound function using PsychoPy (Peirce, 2007) and played in sequences either synchronously (SYNC) or asynchronously (ASYNC). The total number of stimuli was 500 sounds in the main experiment and 250 in the control experiment. Sequences contained standard (80%) and deviant (20%) tones, deviants being either a silence (OMISSION) or a different tone (RARE TONE) (400Hz *vs*. 800Hz, counterbalanced across participants) that appeared every 5 sounds (REGULAR) or between 3 to 7 sounds (uniformly distributed, RANDOM) (Figure 1 top). Both experiments followed the same 3x2 factorial within-subject design (Figure 1 bottom).

**Figure 1.**
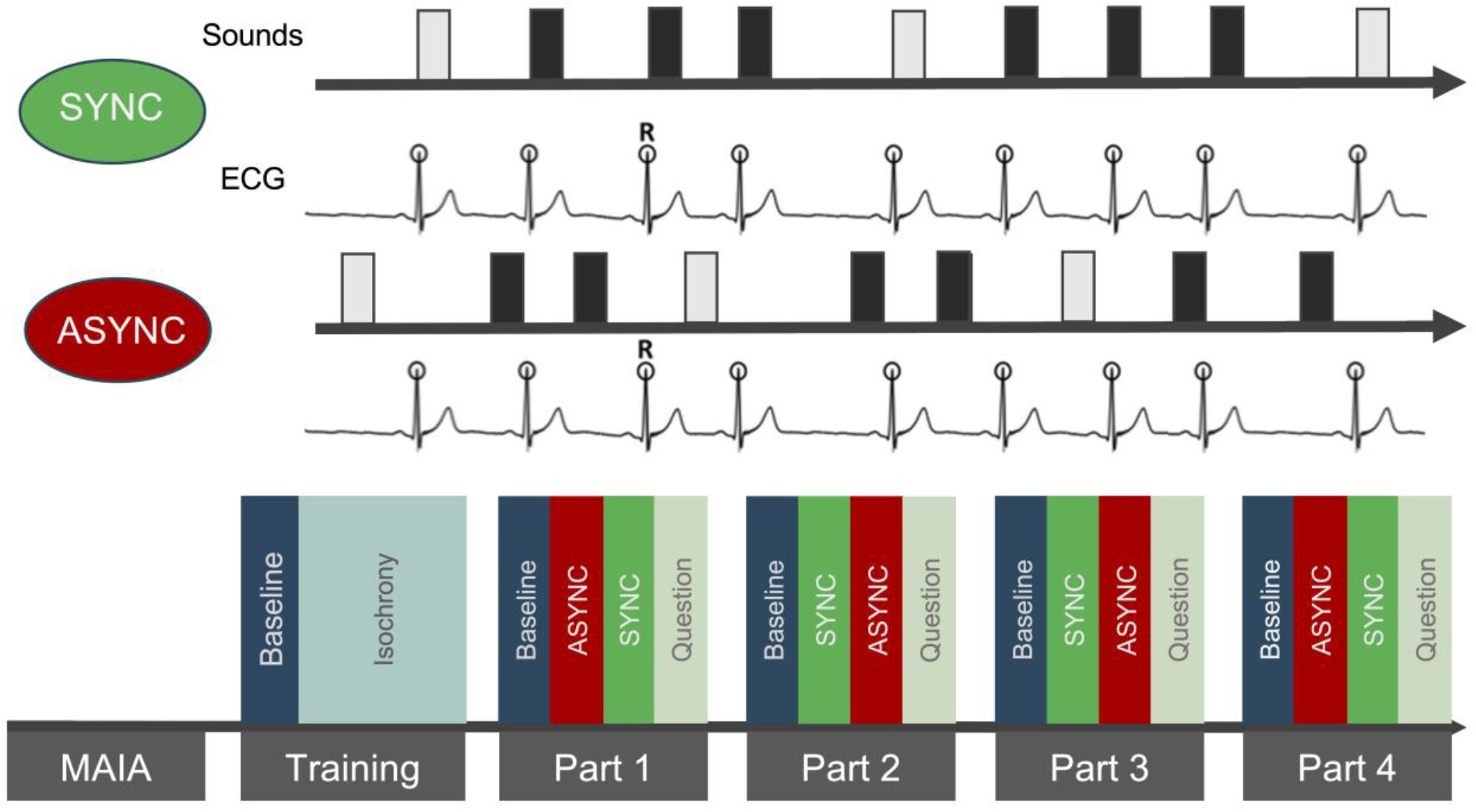
Experimental protocol. The cardio-audio synchrony paradigm was implemented by presenting standard tones (black bar) and deviants (grey bar) either synchronously (SYNC) or asynchronously (ASYNC) to the participants’ heartbeats. In both behavioral and physiological experiments, the same 3x2 factorial within-subject design was used. After a training session, four parts containing a different combination of cardio-audio synchronization, type of deviant and regularity were presented. Interoceptive abilities were assessed before the experiment by administering the FR-MAIA questionnaire and subjective awareness of the cardio-audio synchrony at the end of each part by asking participants a discrimination question which of the SYNC or ASYNC condition was played first.

Participants started by a training phase of a 30s baseline at rest followed by 4 training trials displaying a different combination of type of deviant (OMISSION or RARE TONE) and regularity (RANDOM or REGULAR) presented at a fixed, *i*.*e*., isochronous, sound-to-sound interval matching the mean heart rate of the baseline. The testing phase contained four parts, each presenting a different combination of regularity and deviant type following the same order as training trials (order counterbalanced across participants). For each part, a baseline (10 minutes in the main experiment, 5 minutes in the control experiment) was followed by two sound sequences either synchronous (SYNC) or asynchronous (ASYNC) (order counterbalanced across parts within each participant).

In the main experiment, participants were asked to simply attend to sounds. In the control experiment, participants were required to press a button as soon as they detected a deviant. In both experiments, participants were asked after each part whether the SYNC or ASYNC condition was presented first using a six-point Likert’s confidence scale (1: very confident SYNC to 6: very confident ASYNC). After the experiment, participants were debriefed by asking them whether they noticed a particular pattern in the sound sequences and whether they used any strategy to distinguish between SYNC and ASYNC conditions.

### Synchronisation

Temporal onsets of sounds were defined based on the online analysis of real-time ECG recordings using an open-source Python package (Scheltienne 2022; mscheltienne/cardio-audio-sleep: 0.3.0. Zenodo. https://doi.org/10.5281/zenodo.7057572) (Figure 2). R-peaks were detected online over the last 4 seconds of stream acquired when the signal satisfied the peak detection criteria (height percentage = 97,5%, prominence = 500, width = NONE, interbeat interval = 500ms). When necessary, the detection criteria were individually adapted prior to the experiment.

**Figure 2.**
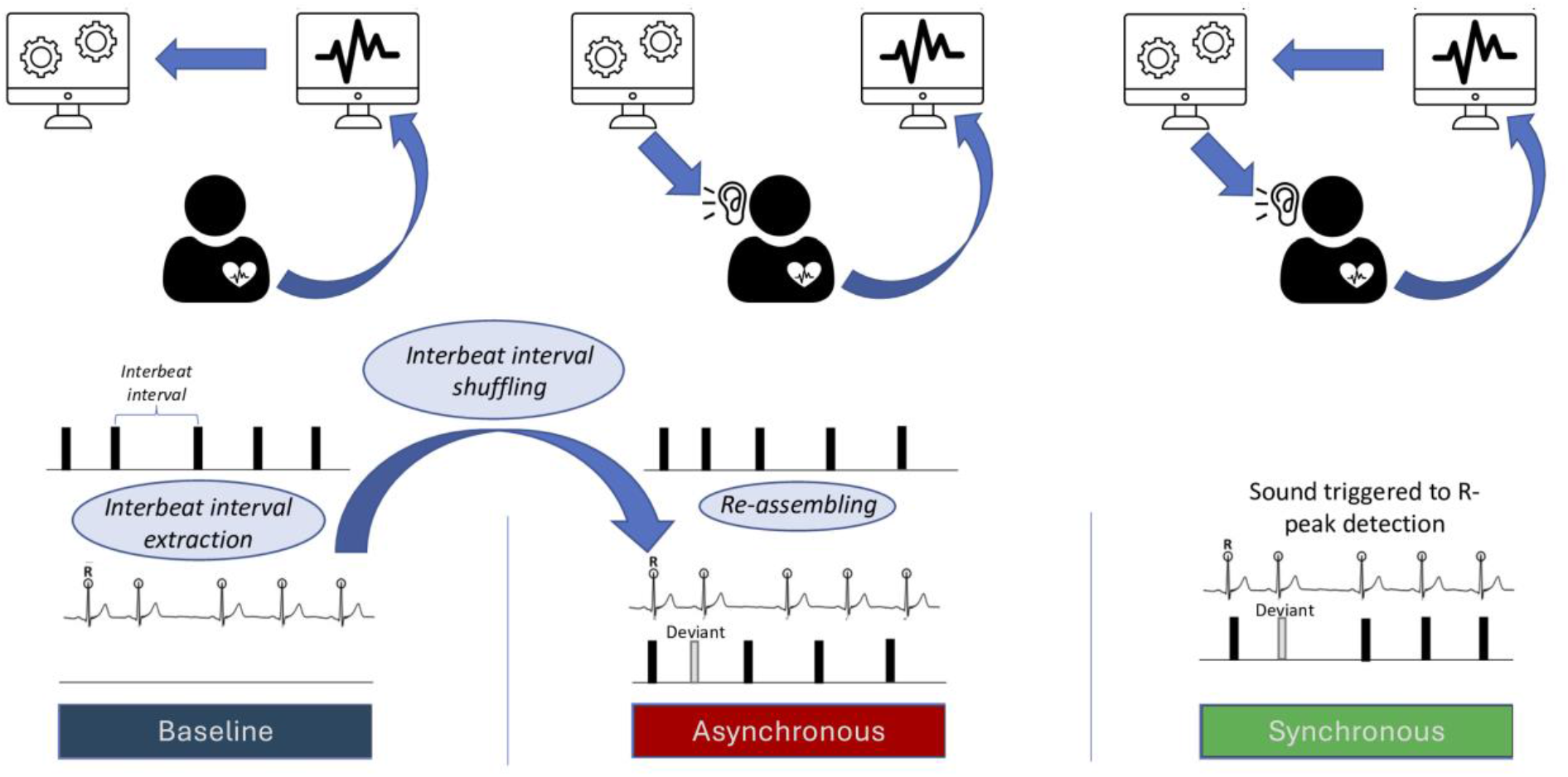
Synchronisation procedure. ECG was continuously recorded and the R-peaks were extracted online. For each part, the obtained interbeat interval sequence during a baseline at rest without any sound stimulation was shuffled and used to generate a pseudo-sequence of sound stimulation. This pseudo-sequence was used in the asynchronous condition to trigger sound stimulation at a frequency matching the distribution of interbeat intervals of the participant. In the synchronous condition, R peaks were extracted online using a closed-loop stimulation algorithm to trigger sound presentations at a short delay (50ms on average) after each heartbeat.

In the SYNC condition, sounds were triggered upon R peak detection with an R-peak to sound onset (RS) delay aimed at 50ms (49.4ms±1.3 [mean±standard error] across participants in the main experiment, and 52.2ms±2.6 in the control study). In the ASYNC condition, interstimulus intervals were determined by randomly shuffling the interbeat intervals of the ECG signals from the baseline trial following Pfeiffer *et* De Lucia (2017).

This resulted in a more variable RS delay (ASYNC: 277.5ms±[277.8, 292] *vs*. SYNC: 90.1ms±[80.6, 99.5] (mean and 95% confidence interval), post-hoc Estimated Marginal Means test: 280ms±2.78 (mean±standard error), p<0.001) and Sound to R-peak (SR) delay for the ASYNC condition (standard variability for ASYNC: 282.0ms±[272.5, 291.4] *vs*. SYNC: 4.99ms±[-2.06, 12], 192ms±2.15, p<0.001) (Figure 3 top).

**Figure 3.**
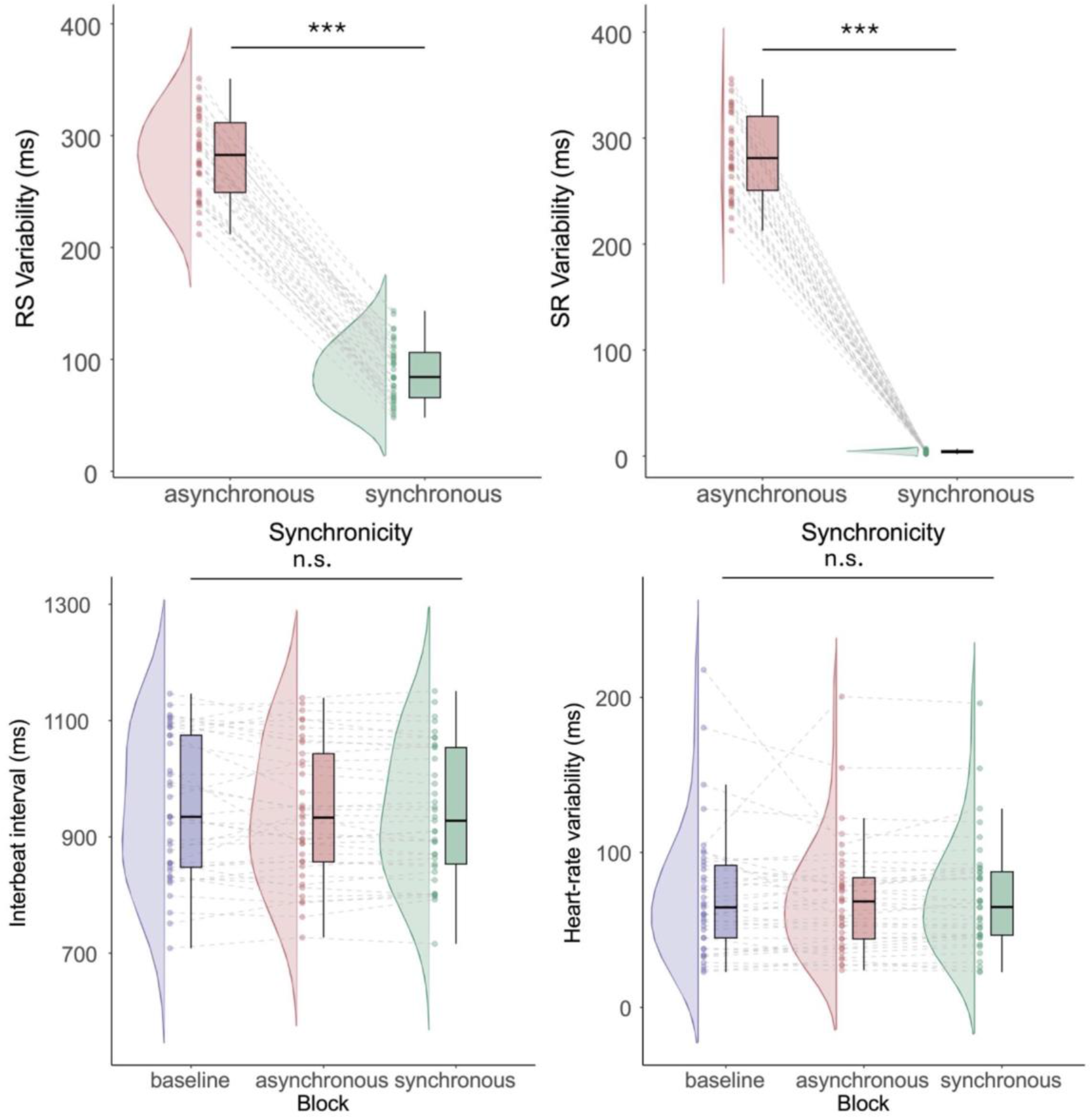
Control analyses. The standard deviation of the interval between the R-peak of the heartbeat (R) and the onset of the deviant (S) was reduced for synchronous over asynchronous conditions before (top left panel) and after (top right panel) the occurrence of the deviant. The average (bottom left panel) and the root-mean square of the standard deviation (bottom right panel) of the interbeat interval distribution were not affected across each type of block over the entire experiment for each participant. Post-hoc Estimated Marginal Means test: ***, p<0.05, n.s., not significant.

We finally confirmed that this procedure did not affect heart rate (interbeat interval for ASYNC: 942ms±[903, 981] *vs*. SYNC: 940ms±[901, 979], post-hoc Estimated Marginal Means test: 1.60ms±5.41, p=0.953) and heart rate variability (root-mean square of standard deviation of interbeat interval for SYNC: 71.3ms±[58.3, 84.2] *vs*. ASYNC: 70.1ms±[57.2, 83.1], post-hoc Estimated Marginal Means test: -2.76ms±3.78, p=0.951) (Figure 3 bottom).

### Analysis

For the main experiment, ECG data was preprocessed offline by applying a 0.5 Hz high-pass Butterworth and powerline filtering (order = 5, powerline = 50 Hz) using the open-source Python neurokit2 package (Makowski et al., 2021). Heartbeats were extracted by detecting R-peaks using the neurokit method and interbeat intervals were computed around deviants’ onset. The outcome variable was the modulation of the heartbeat expressed in the form of interbeat intervals. A linear mixed model was applied with condition as fixed effects and subjects as random effects using the lmer function of the lme4 package in R (Bates et al., 2015). Within-condition differences were assessed by performing post-hoc tests using the Estimated Marginal Means method of the emmeans package (Piaskowski, 2025).

For the control experiment, the outcome variable was the reaction time to detect deviants. Reaction times were obtained for each trial by computing the difference between the onset of the deviant and button press, excluding values three standard deviations above the mean of each participant. A generalized mixed model was applied with condition as fixed effects and subjects as random effects with a gamma distribution and inverse link function. Effects of conditions were assessed by testing the goodness of model fit on the condition tested using likelihood ratio tests. Within-condition differences were assessed by performing post-hoc tests using the Estimated Marginal Means method.

For both experiments, discrimination of synchronous and asynchronous conditions was investigated by scoring correct responses as positive values (*i*.*e*., 0.33 for random, 0.66 for unsure, 1 for certain) and incorrect responses as negative values (0.33 for random, 0.66 for unsure, 1 for certain) and averaging scores for each participant. Response scores were analyzed using non-parametric Wilcoxon tests against 0 to investigate the ability of correctly guessing which block was synchronous and asynchronous. Interoceptive ability was measured as the total score of the MAIA questionnaire (Willem et al., 2022). Kendall correlation tests were used to assess the correlation of response scores with FR-MAIA scores with cardiac responses.

## Results

### Cardiac activity is modulated by cardio-audio synchrony only

Four out of 40 participants were excluded from the analyses of the main experiment due to a technical failure in the sound-cardiac synchronization, resulting in the inclusion of a total of 36 participants (age=23.0±0.6, 18 female). Linear mixed model effects revealed that cardiac responses were modulated after the occurrence of auditory deviants, with a main effect of cardio-audio synchrony (*β*=0.03±[0.02, 0.03], t(1142)=8.46, p<0.001), but not deviant type (*β*=2.0.10^-3^±[-4.5.10^-3^, 8.6.10^-3^], t(1142)=0.61, p=0.54) nor regularity (*β*=7.1.10^-4^±[-5.68.10^-3^, 7.11.10^-3^], t(1142)=0.22, p=0.83) (Figure 4).

**Figure 4.**
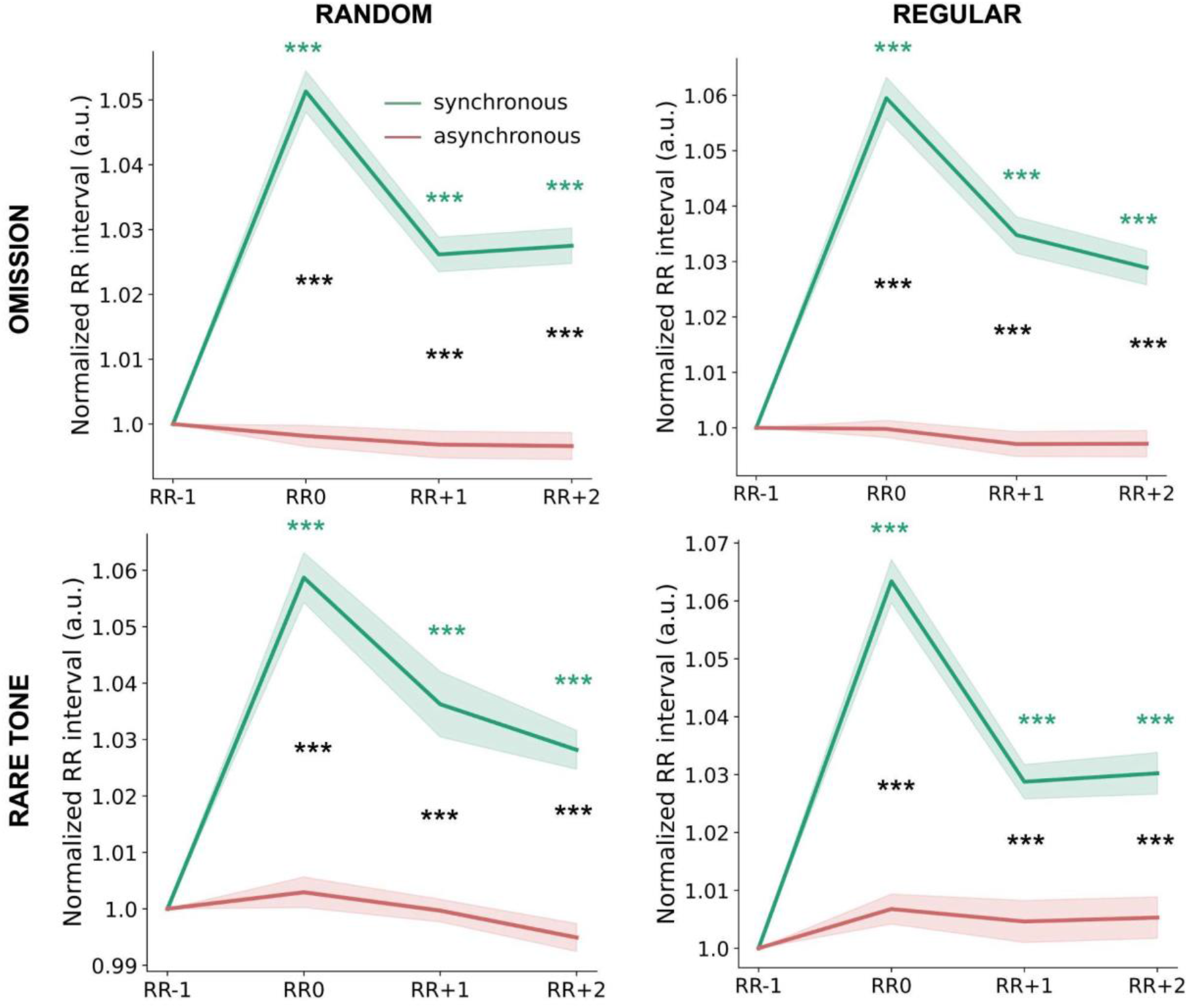
Cardiac slowdown results from synchrony, but not predictability or the type of deviant. The interbeat interval around deviant’s onset (RR0) and after one (RR+1) and two heartbeats (RR+2) are greater compared to the interbeat interval preceding deviant’s onset (RR0) in arbitrary units (a.u) in the synchronous (green), but not in the asynchronous (red) conditions. Cardiac responses are shown for random omission (top left), regular omission (top right), random rare tone (bottom left) and regular rare tone (bottom right). Solid lines represent the mean and shaded areas the standard error to the mean. Post-hoc Estimated Marginal Means test for synchronous (green) and between synchronous and asynchronous conditions (black), ***: p<0.001.

To compare interbeat intervals (RR) across conditions, we normalized RR following deviants’ onsets by the one preceding deviants’ onsets (RR-1). Post-hoc pairwise tests revealed that the heartbeat slowed down after the occurrence of deviants (RR0) for the synchronous, but not the asynchronous condition (RR0 vs RR-1 for synchronous: 0.06±2.4.10^-3^ms, p<0.001; and asynchronous: 0.00±2.4.10^-3^, p=0.87, synchronous *vs*. asynchronous at RR0: 0.06±3.1.10^-3^ms, p<0.001), as well as for the following ones (see Table S1 for a full description of the results).

We also checked that this interpretation of results could not be limited by heart rate variations across conditions. Linear mixed-models confirmed the absence of a significant difference in heart rate across conditions, excluding this confound (see Table S2 for a full description of the results). Overall, our results demonstrate that cardiac activity slow down after deviants’ onset when sounds were synchronized with heartbeats, independent of the type and regularity of deviants.

### Behavioral reaction is influenced by both predictability and deviant’s type

As planned, 10 healthy adults were included in the control experiment (age=24.8 years±2.4; mean±standard error, 5 females). To check how auditory deviations influenced participants at the behavioral level, we investigated how cardio-audio synchrony, regularity and the type of deviant impacted their reaction time to detect deviants. The generalized linear mixed model (GLMM) analysis revealed no evidence of an effect of cardio-audio synchrony (likelihood test ratio with and without synchrony as a co-factor, *χ*2=4.91, p=0.30). Yet, we found a main effect of type of deviant (*β*=0.04s±[0.03,0.05], t(3721)=8.73, p<0.001) and regularity (*β*=0.12s±[0.11, 0.13], t(3721)=25.30, p<0.001), as well as an interaction effect between both deviants and regularity (*β*=-0.07s±[-0.09, -0.06], t(3721)=-11.27, p<0.001) (Figure 5).

**Figure 5.**
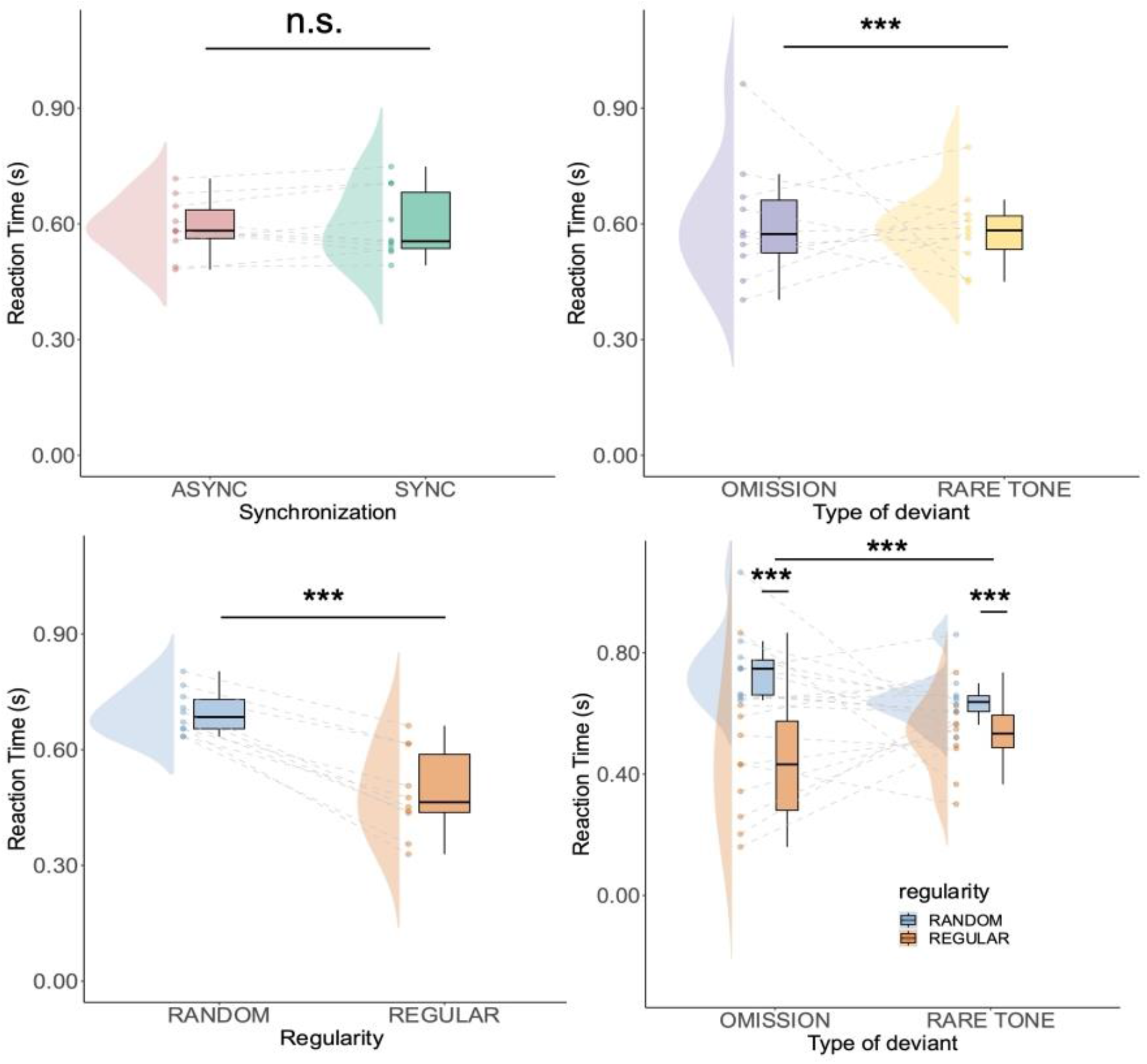
Behavioral responsiveness is influenced by the type of deviant and regularity, but not synchrony. There was a faster detection of regular omissions over random, and of regular tones over random ones. No main effect of synchronicity (top left) were found, yet omissions (top right) and regular tones (bottom left) were detected faster. Data points represent single-subject averages, boxplot shows the median, first and third quartile and extreme values, half violin plot shows the distribution across the population for each condition. Post-hoc Estimated Marginal Means tests, n.s.: non-significant, ***: p<0.001.

Post-hoc pairwise tests further revealed that regular omissions were detected faster than random omissions (difference: -0.30s±0.02, p<0.001), as well as for regular compared to random rare tones (-0.12s±0.01, p<0.001), confirming that expectations of upcoming auditory deviants speed up behavioral reactions. We also found that random omissions were detected more slowly than random rare tones (0.11s±0.01, p<0.001), showing that rare tones are detected faster than omissions whenever they could not be predicted. This relationship was inverted for regular omissions *vs*. rare tones (-0.08s±0.01, p<0.001), showing that whenever predictable, omissions were detected faster than rare tones.

The fact that button presses to regular omissions could be observed even before the occurrence of deviants further suggests that participants anticipate the deviants’ omission in absence of incoming sensory information. These observations suggest that participants actively infer the presence of deviants, showcasing the role of predictive processes in behavioral reactions. Overall, these results confirm that the detection of deviants is influenced by their type and their predictability at the behavioral level, in contrast to the findings obtained at the cardiac level, where cardiac deceleration was observed independent of deviation type.

### Subjective decision reflects cardio-audio synchrony

We finally investigated whether participants detected which sequence was synchronous or asynchronous at the subjective level. Model comparison of generalized mixed models (GGLM) showed no significant differences between the ratings of the main and control experiments (Likelihood-ratio test with and without experiment as a co-factor, *χ*2=0.43, p=0.93). Across both experiments, GLMM analysis revealed that confidence scores accurately predicted correct responses (*β*=0.85±[0.12, 1.57], p=0.022). To account for this interaction in post-hoc tests, we re-scored subjective responses by ordering confident incorrect responses to confident correct responses on a 6-point Likert scale.

Post-hoc tests on response scores confirmed that participants identified cardio-audio synchrony above chance level (non-parametric Wilcoxon-test test: r=0.52, p<0.001 on both experiments). Additionally, response scores did not correlate with MAIA scores across both experiments (Kendall correlation test, tau = 0.050, p-value = 0.625). Further, we found no evidence that cardiac responses in the main experiment correlate with response (Kendall’s tau= 0.002, p-value = 0.924) or MAIA scores (Kendall’s tau = -0.005, p-value = 0.823). Overall, we observed that participants’ ability to identify which sound sequences were synchronized with their heartbeat but could not find a relationship with their individual interoceptive ability nor their cardiac responses to auditory deviants (Figure 6).

**Figure 6.**
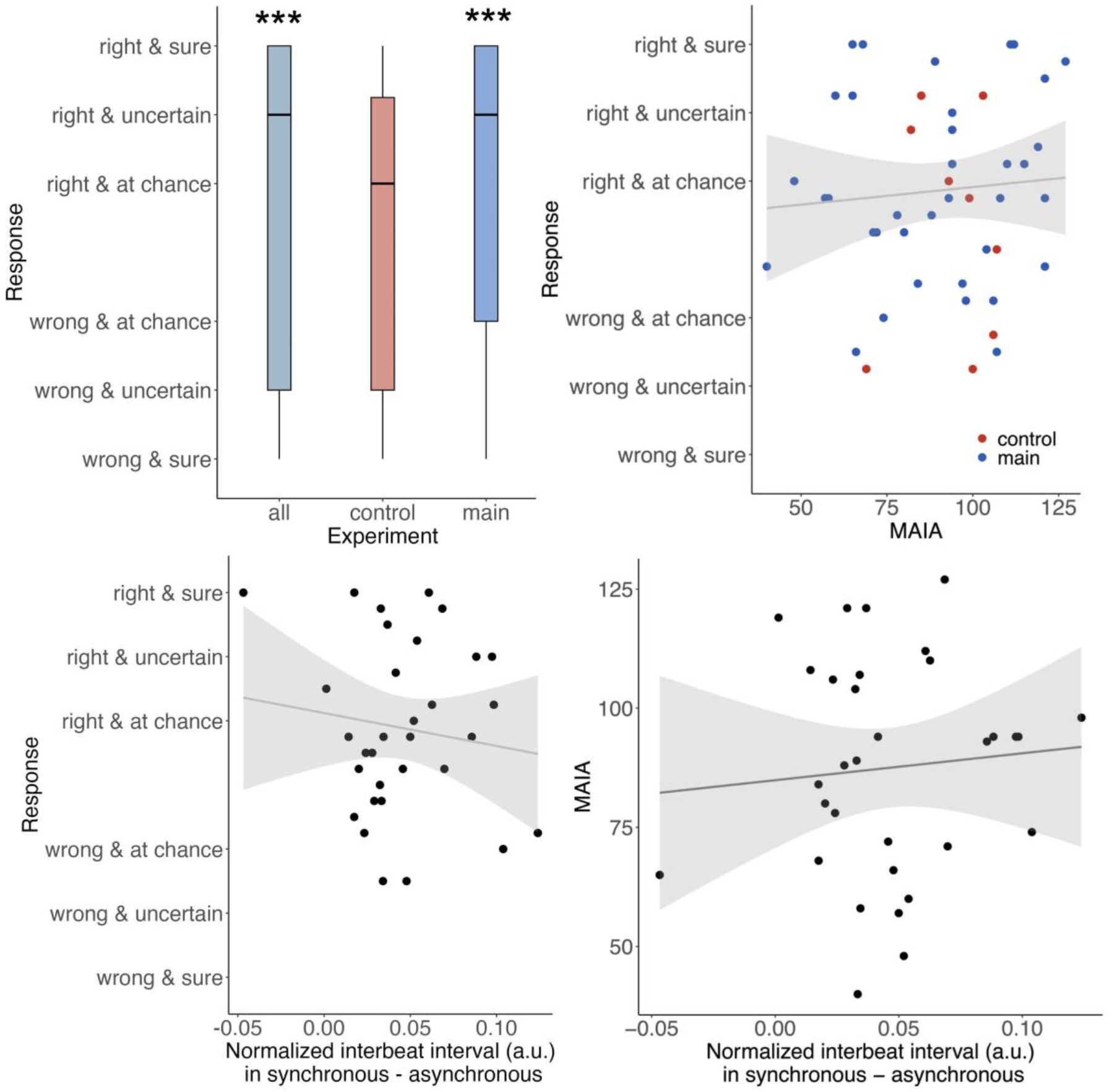
Participants were able to subjectively detect cardio-audio synchrony, without evidence for a systematic relationship with interoceptive abilities and cardiac modulation. Responses were on average significantly correct, scored as positive values (i.e., 0.33 for random, 0.66 for unsure, 1 for certain decision) along with incorrect responses scored as negative values (-0.33 for random, -0.66 for unsure, -1 for certain decision). Boxplots represent the median, first and third quartile and extreme values of responses (top left panel). Data points represent averages of subjects for response and total MAIA scores. Solid line and shaded areas represent the linear regression of the correlation between variables (top right and bottom panel). Wilcoxon rank-sum test, ***: p<0.001

Besides the lack of significant evidence on the source of information used to guide subjective decision, we obtained anecdotical reports from explorative post-experiment interviews that some participants noticed that their breathing patterns influenced the occurrence of sounds, allowing them to correctly guess which block presented sounds synchronously with their heartbeats. This suggests that participants could rely on the visceral coupling between respiratory activity and cardiac rhythm, rather than the direct monitoring of their heartbeat to detect cardio-audio synchrony. Overall, our results demonstrate the ability of participants to detect cardio-audio synchrony at the subjective level without evidence of a systematic relationship with interoceptive abilities and cardiac modulation.

## Discussion

Our study investigated cardiac and behavioral responses to heartbeat-locked auditory deviations to inform the mechanistic underpinnings of interoceptive-exteroceptive integration according to the theoretical accounts of cardiac surprise, active inference, and dynamical coupling. To this aim, we tested the role of sensory and predictive processes by varying the type of deviation (omissions vs. rare tones) and their predictability (random vs regular). We observed that heartbeat-locked auditory deviations slow down cardiac activity independent of their type and regularity. In contrast, participants’ reaction times to deviant sounds were sensitive to the type of deviation and stimulus predictability. Finally, despite participants’ ability to detect cardio-audio synchrony, their recognition scores did not correlate with interoceptive ability nor cardiac responses. Overall, our results suggest that cardiac, behavioral and subjective responses in cardio-audio synchrony might operate, at least partially, through different mechanisms.

Our results generalize previous findings that cardiac activity transiently decelerated after omissions when sounds were synchronized to heartbeats, showing evidence for interoceptive-exteroceptive integration (Banellis & Cruse, 2020; Pelentritou et al., 2024, 2025). This cardiac deceleration can be interpreted as resulting from the action of a parasympathetic brake later removed upon the reinstatement of standard tones, allowing heart rate to return to baseline levels (Skora et al., 2022). More generally, cardiac slowdown has been associated with error-related processing (Skora et al., 2022) and the detection of threatening stimuli (Battaglia et al., 2024; Livermore et al., 2021), as exemplified by freezing responses that reduce the internal noise associated with heartbeats and favor the processing of incoming signals (Jennings et al., 1992; Roelofs, 2017; Skora et al., 2022). Here, we showed that cardiac slowdown observed when sounds are synchronized with heartbeats was independent from the deviants’ type and predictability, going against our initial expectations based on our pre-registered hypotheses.

We consider that this set of results suggests that cardiac responses during cardio-audio synchrony do not rely on cardiac surprise, dynamical coupling or active inference as previously proposed (Banellis & Cruse, 2020; Pelentritou et al., 2024; Pfeiffer & De Lucia, 2017; Banellis & Cruse, 2021). Previous works demonstrated that the synchronization of stimuli with heartbeats caused them to be associated with the processing of the self (Sel et al., 2016; Suzuki et al., 2013). In line with this interpretation, previous findings showed that heartbeat evoked potentials, *i*.*e*., neural responses to internally generated cardiac signals, were enhanced following omissions of sounds synchronized with heartbeats (Banellis & Cruse, 2020; Pfeiffer & De Lucia, 2017). The cardiac slowdown to auditory deviations may, thus, be a response to salient events associated with the self. Noteworthy, this implies that cardiac responses to cardio-audio synchrony could be used to probe self-related processes without relying on behavioral or subjective reports, which is particularly relevant in conditions of limited behavioral responsiveness or awareness, such as disorders of consciousness, anesthesia, sleep or psychedelic states.

Contrary to our predictions, we could not obtain evidence that reaction times of deviants were modulated by cardio-audio synchrony, *i*.*e*., synchronous *vs*. asynchronous presentation of sounds to participants’ heartbeats. These results go along with previous findings demonstrating that cardio-audio synchrony did not affect auditory detection tasks at the behavioral level (Banellis & Cruse, 2020; Delfini & Campos, 1972). We interpret the interaction between participants being quicker to detect regular rare tones (over omissions) and random omissions (over rare tones) as evidence that participants anticipate the presence of deviants whenever they are predictable, especially when inputs are absent in the case of omission. Overall, our findings suggest that participants actively infer the occurrence of salient auditory deviations in their behavioral reactions without significant evidence of using the exteroceptive-interoceptive integration of cardiac signals during cardio-audio synchrony.

Nevertheless, we found, in line of our prediction, that participants discriminated which condition was synchronous or asynchronous, following previous evidence of subjective awareness of interoceptive-exteroceptive integration using a different cardio-audio synchronization procedure (Banellis & Cruse, 2020; Delfini & Campos, 1972). Subjective responses did not correlate with interoceptive abilities nor showed a relationship to cardiac activity. Together, these findings suggest that cardiac responses to deviants during cardio-audio synchrony might operate independently from subjective awareness. This interpretation aligns with evidence of a cardiac slowdown after heartbeat-locked auditory deviations in participants with reduced awareness in deep non-rapid eye movement (NREM) sleep (Pelentritou et al., 2024) and coma (Pelentritou et al., 2025). Anecdotal reports that breathing patterns can guide participants’ detection of cardio-audio synchrony further suggest different mechanisms by which conscious subjects discriminate whether external signals are synchronized with their internal activity. For example, rather than using cardiac signals to directly track interoceptive-exteroceptive integration, participants could resort to the dynamical coupling of respiratory activity with external sounds to do so.

We acknowledge limitations of this paper by considering further analyses that can be carried out with this paradigm. The study of reaction times using drift-diffusion modeling could refine our understanding of how priors and evidence accumulation are involved in the detection of auditory deviations (Ratcliff et al., 2016). Signal detection theory could be used to assess the origin of subjective detection of cardio-audio synchrony in terms of priors and sensitivity (Macmillan, 2002). The investigation of EEG responses to heartbeats and deviants could address how physiological and sensory signals are integrated at the cortical level (Pfeiffer *et* De Lucia, 2017; Pelentritou et al., 2025). Electrodermal and pupil activity as physiological correlates of arousal can be used to investigate affective markers of auditory mismatch detection to test whether they depend upon the type of deviations and their predictability during cardio-audio synchrony (Kamp & Donchin, 2015; Chiarion et al., 2025). Finally, investigating the modulation of cardiac activity by respiratory patterns could deepen our understanding of the dynamical coupling of visceral signals that may guide subjective detection of cardio-audio synchrony (Schultz et al., 2013).

To conclude, this work provides a comprehensive investigation of the cardiac correlates of interoceptive-exteroceptive integration during cardio-audio synchrony. Our demonstration that heartbeat-locked auditory deviations slow cardiac activity down irrespectively of their type and predictability coincides with no models from the previous literature, leading us to propose a novel interpretation of cardiac responses to cardio-audio synchrony as an implicit marker of self-referential information processing. Finally, the dissociation observed between cardiac, behavioral, and subjective responses suggest that different mechanisms underly these levels. Future investigation of the cardio-audio synchrony could test this hypothesis with original pre-registered predictions and experimental designs to elucidate the mechanisms of interoceptive-exteroceptive integration of cardiac and auditory signals in wakeful and low-arousal states.

## Acknowledgments

We are grateful to Mathieu Scheltienne and the Human Neuroscience Platform for sharing the code supporting the cardio-audio synchronization procedure and their technical support.

## Data availability statement

The code and dataframes supporting this article are available on the OSF page: doi.org/10.17605/OSF.IO/6FVUW. Raw data can be requested to the corresponding author and will be openly accessible with the publication of the article.

## Author contributions

**Matthieu Koroma**: Conceptualization; investigation; writing – original draft; methodology; validation; visualization; writing – review and editing; project administration; formal analysis; software; data curation; resources; funding acquisition; supervision.

**Kevin Nguy**: Conceptualization; investigation; writing – original draft; methodology; validation; visualization; writing – review and editing; formal analysis; software; data curation. **Andria Pelentritou**: Conceptualization; methodology; validation; writing – review and editing; resources.

**Marzia De Lucia**: Conceptualization; Methodology; validation; visualization; writing –review and editing; resources; supervision.

**Athena Demertzi**: Supervision; conceptualization; investigation; validation; visualization; writing – review and editing; formal analysis; project administration; resources; data curation; methodology; writing – original draft; funding acquisition.

