## Supplementary Information for "Heartbeat-locked auditory deviations slow down cardiac activity"

*Table S1* ***Full details of the cardiac responses at different delays after sound deviants***. *The normalized interbeat intervals between the occurrence before and after the presentation of deviants (RR0), the first and second heartbeat following sound occurrence (RR+1) and the second and third heartbeat after sound occurrence (RR+2) for each condition are detailed. The mean and confidence interval for synchronous condition (SYNC) and asynchronous condition (ASYNC) are reported, as well as the mean and standard error to the mean interval difference between synchronous and asynchronous conditions (Diff). Statistical tests are Wilcoxon-rank sum test between synchronous and asynchronous condition, ***: p<0.001*

| **Type of deviations** | | Omission | | Rare tone | |
|---|---|---|---|---|---|
| **Predictions** | | Random | Regular | Random | Regular |
| **RR0** | SYNC | 1.05±[1.04,1.05] | 1.06±[1.05,1.07] | 1.06±[1.05,1.07] | 1.06±[1.05,1.07] |
| | ASYNC | 1.00±[0.99,1.00] | 1.00±[0.99,1.01] | 1.00±[0.99,1.01] | 1.01±[1.00,1.02] |
| | Diff | 0.049±0.004*** | 0.056±0.005*** | 0.053±0.007*** | 0.050±0.005*** |
| **RR +1** | SYNC | 1.04± [1.03,1.05] | 1.04±[1.03,1.05] | 1.04±[1.03,1.05] | 1.04±[1.03,1.05] |
| | ASYNC | 1.00±[0.99,1.01] | 1.00±[0.99,1.01] | 1.00±[0.99,1.01] | 1.01±[1.00,1.02] |
| | Diff | 0.037±0.003*** | 0.043±0.003*** | 0.042±0.004*** | 0.034±0.003*** |
| **RR +2** | SYNC | 1.02±[1.02,1.03] | 1.03±[1.02,1.03] | 1.03±[1.01,1.04] | 1.02±[0.99,1.02] |
| | ASYNC | 1.03±[0.99,1.00] | 1.00±[0.99,1.01] | 1.00±[0.98,1.01] | 1.01±[1.01,1.03] |
| | Diff | 0.026±0.004*** | 0.029±0.003*** | 0.031±0.007*** | 0.018±0.005*** |

*Table S2* ***Full details of heart rate across the different conditions****. Heart rates computed according to interbeat intervals in ms. The mean and confidence interval for synchronous condition (SYNC) and asynchronous condition (ASYNC) are detailed, as well as the mean and standard error to the mean of the difference between synchronous and asynchronous conditions (Diff). Statistical tests are Wilcoxon-rank sum tests between synchronous and asynchronous conditions, n.s. : not significant*

| **Type of deviations** | | Omission | | Rare tone | |
|---|---|---|---|---|---|
| **Predictions** | | Random | Regular | Random | Regular |
| HR | SYNC | 937±[896, 976] | 1.06±[1.05,994] | 939±[899,979] | 935±[895, 976] |
| | ASYNC | 938±[898, 979] | 953±[913, 992] | 938±[897,979] | 936±[895, 977] |
| | Diff | -2.34±4.86 n.s. | -1.80±4.88 n.s. | -0.78±4.35 n.s. | -1.48±4.76 n.s. |
